# A new species of *Mesopolobus* Westwood (Hymenoptera, Pteromalidae) from black locust crops

**DOI:** 10.1101/2020.05.01.072140

**Authors:** Zoltán László, K. Tímea Lakatos, Avar-Lehel Dénes

## Abstract

A new species of the genus *Mesopolobus* Westwood, *Mesopolobus robiniae* sp. n., is described and illustrated from east-central Europe (Romania and Hungary). The species was reared from black locust (*Robinia pseudoacacia*) seedpod samples, where it most likely parasitizes the black locust’s seed predator *Bruchophagus robiniae* Zerova, 1970. Here we present the new species and report on its ecological relationships within the European seed predator community of black locust. We also give details regarding type material and type locality, a detailed description with images, a differential diagnosis of the new species, and a modification to the identification key published by Graham (1969), that distinguishes this new species from closely related species. In addition, we provide information on the distribution, biology and results of barcoding analysis. We also provide the DNA sequence data to complement the morphological taxonomy.

## Introduction

In the last century the black locust (*Robinia pseudoacacia* L.) became a characteristic component feature of the Central and Eastern European landscape (Vítková *et al.*, 2017). Its positive economic, but negative environmental impacts led to conflicts between nature conservationists, forestry workers, urban planning experts, beekeepers and the public (Benesperi *et al.*, 2012; Dickie *et al*., 2014; Sádlo *et al*., 2017). As current legislation will determine the future distribution of black locust, we need detailed knowledge, not only from the viewpoint of the forestry and economy, but also from the viewpoint of the species potential associates, like herbivorous insects and their community (Kleinbauer *et al.*, 2010).

The invasive history of black locust follows the characteristic pathway of introduced crops with an initial phase when presumably several independent introductions occurred from North America, which ceased for a long period, then were followed by frequent plantings and a rapid invasion in the wild, resulting in its widespread distribution of today (DAISIE, 2009). The invasion of black locust in Central and Eastern Europe was facilitated by extensive plantings, due to the wood’s long-term quality, resistance to insects and fungi, rapid growth, easy propagation, and ability to stabilize soils (Vítková *et al.*, 2017).

When replacing native vegetation, the black locust reduces local biodiversity (Hanzelka & Reif, 2015). Endangered light-demanding plants and invertebrates are threatened by its appearance through reducing light to plants growing beneath the canopy and above the forest floor, and changing the microclimate and soil quality (Lazzaro *et al.*, 2018). These impacts can have effects throughout the food chain, by depriving birds of their insect prey, which depend on the plants that have been wiped out by the black locust (Hanzelka & Reif, 2015). One of the central problems regarding black locust colonization is its capacity to rapidly increase soil nutrient concentration and to alter soil chemical properties which conditions then facilitate invasion by other non-native nitrophilous plant species (Enescu & Dănescu, 2013).

Because of the wide distribution and negative environmental impacts of black locust an important management tool for invasive populations is limiting their propagation (Redei *et al.*, 2001). The crops of black locust are attacked by several pests, among which the seed predators can have a major impact (Zerova, 1970; Perju, 1998). *Bruchophagus robiniae* Zerova, 1970 (Hymenoptera: Chalcidoidea: Eurytomidae) is a pre-dispersal seed predator, monophagous on black locust seeds (Perju, 1998). As with other *Bruchophagus* species, each *B. robiniae* individual feeds and develops inside one infested seed, so each seed wasp consumes only one seed and each seed houses only one seed wasp (Lakatos *et al.*, 2018). This seed predator has a component community comprising several species, including parasitoid wasps (Lakatos *et al.*, 2016; 2018).

One member of the *Bruchophagus robiniae* - black locust seed predator community belongs to the genus *Mesopolobus* Westwood, 1833, a group of parasitoid wasps in Pteromalidae (Hymenoptera: Chalcidoidea) containing more than 120 described species (Noyes, 2020), over 60 of which are present in Europe (http://www.fauna-eu.org/). Species of this genus have a wide host range, although several species are known to be host-specific on galls (Diptera: Cecidomyiidae, Hymenoptera: Cynipidae), bark beetles (Scolytinae), seed predators (Lepidoptera, Coleoptera: Curculionidae, Bruchinae, Hymenoptera: Eurytomidae), etc. (Bouček & Rasplus, 1991; Noyes, 2020). Several Eurytomidae species, such as *Bruchophagus gibbus* (Boheman, 1836) has *Mesopolobus* sp. parasitoids (Noyes, 2020), and several *Mesopolobus* species are parasitoids of pests on oilseed rape (*Brassica napus* L.), on alfalfa (*Medicago sativa* L.), or on Norway spruce (*Picea abies* L.) (Noyes, 2020).

*Mesopolobus* is a taxonomically complex genus, considering the high number of species belonging to this genus. European *Mesopolobus* were revised by Hans von Rosen (von Rosen, 1958; 1959; 1960; 1961), who synonymized species from multiple genera (*Amblymerus* Walker, 1834; *Eutelus* Walker, 1834; *Platyterma* Walker, 1834) under *Mesopolobus.* The last revision of the *Mesopolobus* genus for the Western Palearctic was written by Graham (1969). Later studies dealing with *Mesopolobus* parasitoids of certain host groups have provided further clarifications of species synonymies, notably parasitoids of gall inducing Cynipidae on *Quercus* sp. (Askew, 1961; Aldrey, 1983; Pujade-Villar, 1993) and of seed weevils associated with Brassicaceae (Baur *et al*., 2007). Since the latest generic revision (Graham, 1969) several new species have been described from Asia (e.g. Narendran *et al*., 2011; Xiao *et al*., 2016) and North-America (e.g. Doganlar, 1979).

The identification of *Mesopolobus* species emerged from black locust crops was based on the most detailed identification key up to date provided by Graham (1969). Using Graham’s keys a number of characters (fore wing marginal vein length ratio to stigmal vein length, number of anelli and funicular segments, position of toruli to anterior margin of clypeus and to median ocellus, pilosity of the basal cell of fore wing, position of hypopygium tip along the gaster) led us to key couplet 16 (page 643), where based on two character combinations, namely gaster length ratio to head plus thorax length and gaster breadth, which did not match our specimens. This suggested the specimens reared from the black locust pods were not represented in the keys, and were likely undescribed. We thus studied several *Mesopolobus* species represented in the keys and compared them morphometrically to the *Mesopolobus* females emerged from black locust seed pods. This approach provides robust insight into *Mesopolobus* morphology, which may play a major role in resolving the species delimitations in biocontrol studies. Complementing the morphometric study, we also analyzed *mt*COI sequences of the emerged *Mesopolobus* females from black locust pods, and compared them to the available *mt*COI sequences from the BOLD System and NCBI databases.

Our objectives were the following: i) to identify those morphometric characters that give the best discrimination of the females emerged from black locust seedpods from other *Mesopolobus* species. ii) to calculate the genetic distance values between the *mt*COI sequence of the females emerged from black locust and the other *Mesopolobus* species. iii) to describe the species of the females emerged from black locust seedpods.

## Materials and methods

To gather information about black locust seedpod insect inhabitants we collected seedpod samples in black locust plantations and patches for four years, in the early spring of 2009 in Romania and between 2013-2015 in Romania and Hungary (Table 1). Samples were placed in plastic cups, containing 20-100 seedpods and covered with punched plastic wrap. Samples were kept in a covered balcony with a temperature and humidity close to outdoors at Babe◻-Bolyai University (Cluj-Napoca, Romania) and at University of Debrecen (Debrecen, Hungary). Emerged individuals were monitored and collected monthly from seedpod samples for a year, and stored in 70% ethanol. The dominant emerging species were the seed predator of black locust seeds, *Bruchophagus robiniae*, and its parasitoid, the undescribed *Mesopolobus* species (Lakatos *et al*., 2018).

**Table 1.** Collection dates, location and number of *M. robiniae* sp. n. individuals. BH: Bihor County, Romania, CJ: Cluj County, Romania, HB: Hajdú-Bihar County, Hungary.

| Collection date | Location | Northing | Easting | N | N male | N female |
| --- | --- | --- | --- | --- | --- | --- |
| 2009 | BH | 47.098202 | 21.975355 | 9 | 7 | 2 |
| 2009 | CJ | 46.827366 | 23.629258 | 44 | 16 | 28 |
| 2013 | BH | 47.098202 | 21.975355 | 4 | 0 | 4 |
| 2013 | CJ | 46.801433 | 23.611995 | 26 | 12 | 14 |
| 2014 | BH | 47.098202 | 21.975355 | 69 | 35 | 34 |
| 2014 | CJ | 46.801433 | 23.611995 | 104 | 62 | 42 |
| 2014 | HB | 47.554773 | 21.591610 | 58 | 48 | 10 |
| 2015 | BH | 47.098202 | 21.975355 | 16 | 8 | 8 |
| 2015 | CJ | 46.801433 | 23.611995 | 23 | 13 | 10 |
| 2015 | HB | 47.554773 | 21.591610 | 30 | 14 | 16 |

Identification and description of the emerged *Mesopolobus* species have been made under an Olympus SZ51 binocular microscope, with an 80X magnification and LED lighting. Images were produced by a Canon EOS 600D and a Canon EF 100mm f/2.8 USM Macro Lens. Morphological nomenclature follows Graham (1969). The provided identification key is modified from the keys to genus *Mesopolobus* of Graham (1969). Type material is deposited in the Museum of Zoology, Babeş-Bolyai University, Cluj-Napoca (MZBBU). Specimen identification codes: holotype–MZBBU HYM000011; 14 paratypes–MZBBU HYM000012-25, measured female specimens: HYM000026-39.

For morphological comparison several specimens were loaned from different museums. The specimens of *Mesopolobus amaenus* (Walker, 1834), *M. apicalis* (syn. *thomsonii*) (Thompson, 1878), *M. aspilus* (syn. *elongates*) (Walker, 1835), *M. diffinis* (Walker, 1834), *M. dubius* (Walker, 1834), *M. fasciiventris* Westwood, 1833, *M. semiclavatus* (Ratzeburg, 1848) and *M. typographi* (Ruschka, 1924) were loaned from the Hungarian Museum of Natural History, Budapest, Hungary (HMNH). Specimens of *M. verditer* (Norton, 1869), *M. mediterraneus* (Mayr, 1903), *M. tibialis* (Westwood, 1833) and *M. xanthocerus* (Thomson, 1878) were loaned from the British Natural History Museum. One female specimen of *M. longicollis* Graham, 1969 was measured using ImageJ from photographs of the type provided by Oxford University Museum of Natural History.

We measured 19 morphometric variables, corresponding to those used in the taxonomy of Pteromalidae for calculating typically used ratios (e.g. Graham, 1969) (Table 2), on a total of 55 dry-mounted *Mesopolobus* females belonging to the above-named species (Supplementary Material: Table S1). Measurements were made with an Olympus SZ51 stereo microscope (objective: 110AL2X; eyepiece: WHSZ10X) under 60× and 80× magnification using a calibrated eye-piece micrometer (2.5 mm subdivided into 100 units). For all measurements we ensured that the points of reference were equidistant from the objective of the microscope.

**Table 2.** The selected morphometric characters of female *Mesopolobus* specimens, abbreviations used, and their description.

| Abbreviation | Character name | Definition | Magnification |
| --- | --- | --- | --- |
| clv.b | Clava breadth | Greatest breadth of antennal clava, lateral view | 80× |
| clv.l | Clava length | Length of antennal clava, lateral view | 80× |
| hea.b | Head breadth | Greatest breadth of head, dorsal view | 60× |
| hea.l | Head length | Head length in dorsal view (Graham, 1969), distance between anterior and posterior margin of the head, measured laterally | 60× |
| ltg.b | Seventh gastral tergite breadth | Greatest breadth of the seventh gastral tergite, greatest distance between the outermost lateral edges of the seventh gastral tergite | 80× |
| ltg.l | Seventh gastral tergite length | Greatest length of the seventh gastral tergite, greatest distance between the anterior and posterior edges of the seventh gastral tergite | 80× |
| mv.l | Marginal vein | Length of marginal vein, distance between the point at which the submarginal vein touches the leading edge of the wing and the point at which stigmal vein and postmarginal vein unite (Graham, 1969) | 80× |
| msc.b | Mesoscutum breadth | Greatest breadth of mesoscutum just in front of level of tegula, dorsal view | 80× |
| msc.l | Mesoscutum length | Length of mesoscutum along median line from posterior edge of pronotum to posterior edge of mesoscutum, dorsal view | 80× |
| mss.l | Mesosoma length | Length of mesosoma along median line from anterior edge of pronotum collar to posterior edge of nucha, dorsal view | 60× |
| ool.l | OOL | Shortest distance between posterior ocellus and eye margin, dorsal view (Graham, 1969) | 80× |
| pcl.l | Pronotal collar length | Length of pronotal collar along the median line from the edge between neck and pronotal collar to anterior edge of mesoscutum, dorsal view | 80× |
| ped.b | Pedicellus breadth | Greatest breadth of pedicel, dorsal view | 80× |
| ped.l | Pedicellus length | Greatest length of pedicel, dorsal view | 80× |
| pol.l | POL | Shortest distance between posterior ocelli, dorsal view (Graham, 1969) | 80× |
| ppd.l | Propodeum length | Length of propodeum measured along median line from anterior edge to posterior edge of nucha, dorsal view | 80× |
| sct.b | Scutellum breadth | Greatest breadth of the scutellum, greatest distance between the outermost lateral edges of the scutellum | 80× |
| sct.l | Scutellum length | Length of scutellum along median line from posterior edge of mesoscutum to posterior edge of scutellum, dorsal view | 80× |
| stv.l | Stigmal vein | Length of stigmal vein, distance between the point at which stigmal vein and postmarginal vein unite apically, and the distal end of the stigma (Graham, 1969) | 80× |

Body ratios of *Mesopolobus* female specimens were analyzed using the Multivariate Ratio Analysis (MRA) tool (Baur & Leuenberger, 2011). Variation structure of *Mesopolobus* specimens was analyzed by PCA in shape space to identify the principal components accounting for the variation. For the visualization of each character’s contribution we used PCA ratio spectrum. Body ratios with best discriminant power were determined using the LDA ratio extractor (Baur & Leuenberger, 2011). Analyses were made with R statistical software version 3.6.3 (R Core Team, 2020).

Genomic DNA was extracted from three individuals using DNeasy Blood and Tissue kits (Qiagen Inc., Valencia, CA), following the protocol provided by the manufacturer. Mitochondrial cytochrome c oxidase subunit I (COI) sequences were amplified using the standard LCO1490 and HCO2198 primer pair (Folmer *et al*., 1994) in a 50 µl reaction volume at a 45°C annealing temperature. PCR products were purified with the Wizard SV Gel and PCR Clean–Up System (Promega, USA) and sent for sequencing to Macrogen Inc. (Korea).

Sequences were downloaded and verified with the Basic Local Alignment Search Tool (BLAST) (Johnson *et al.*, 2008). Further, sequences for all available *Mesopolobus* species were also downloaded from the NCBI database and the BOLD System (for reference numbers see Figure 2). The sequences were aligned using a Clustal W algorithm (Thompson *et al.*, 1994) in BioEdit (Hall, 1999). A phylogenetic tree was inferred in MrBayes (Ronquist *et al*., 2012), assuming a GTR+G+I model. Interspecific *p*-distances were calculated in MEGA X (Kumar *et al*., 2018).

## Results

### Multivariate Ratio Analysis of variation in body size and shape

We first performed a series of shape PCAs on all specimens based on 19 morphometric characters. We identified principal components contributing to morphometric variation of all *Mesopolobus* females without prior species-determination by applying the PCA in isometry free shape space function to all specimens as a single group. Then we applied the PCA in isometry free shape space only to the group of females which were closest to those emerged from black locust seedpods. When we included all females in shape space, PC1 and PC2 accounted for 58% of the variation of the entire sampled population. When analyzing only those species pairs which were closest to our target group in shape space, PC1 and PC2 accounted for 61% and 74% of the variation respectively. The first principal components are congruent with the separation of species, although a clear cut between the clusters could not be established (Figure 1). On the first scatterplot (Figure 1a) only five species (*M. amaenus*, *M. verditer*, *M. sericeus*, *M. typographi* and *Mesopolobus* sp. n.) showed a clear separation from the rest, but because of the overplotting with *Mesopolobus* sp. n we also retained *M. fasciiventris* for further analyses. On the other two scatterplots (Figure 1b and 1c) the selected species show almost clear separations on PC1, while on PC2 are overlapping.

**Figure 1.**
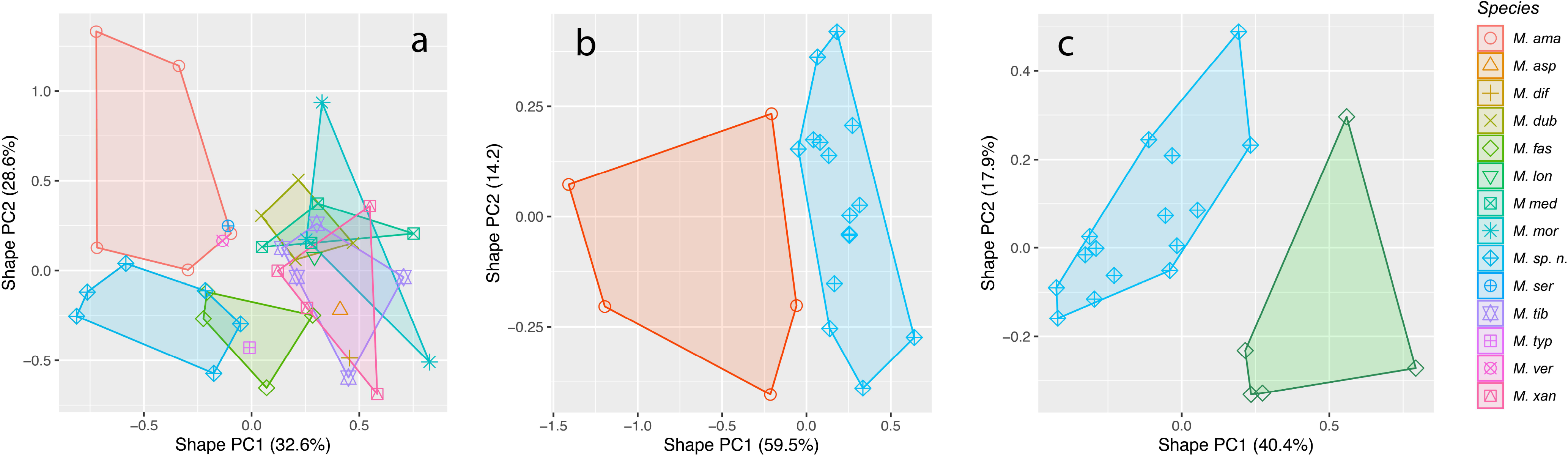
Scatterplot of first against second shape PC based on 19 morphometric variables of females of a) all 14 *Mesopolobus* species b) *M. amaenus* and *Mesopolobus* sp. n emerged from black locust seedpods c) *M. fasciiventris* and *Mesopolobus* sp. n. emerged from black locust seedpods. The variance explained by each shape PC is given in parentheses.

The PCA ratio spectrum for the species pair *M. amaenus* and *Mesopolobus* sp. n. (Figure 2a) identified ltg.l at the extreme high end, and stv.l at the extreme low end of the spectrum. These characters were also found to contribute to species discrimination. The allometry ratio spectrum for the first species pair was dominated almost by the same ratio, stv.l and ltg.l (Figure. 2b), which is also the most important ratio concerning the first shape PC which shows to be the most allometric one. The PCA ratio spectrum for the species pair *M. fasciiventris* and *Mesopolobus* sp. n. (Figure 2c) identified pcl.l at the extreme high end, while stv.l at the extreme low end of the spectrum. These characters, except for stv.l, were found to contribute to species discrimination. The allometry ratio spectrum for the second species pair was dominated almost by the same ratio, pcl.l and ltg.b (Figure. 2d), that is not the most important ratio concerning the first shape PC.

**Figure 2.**
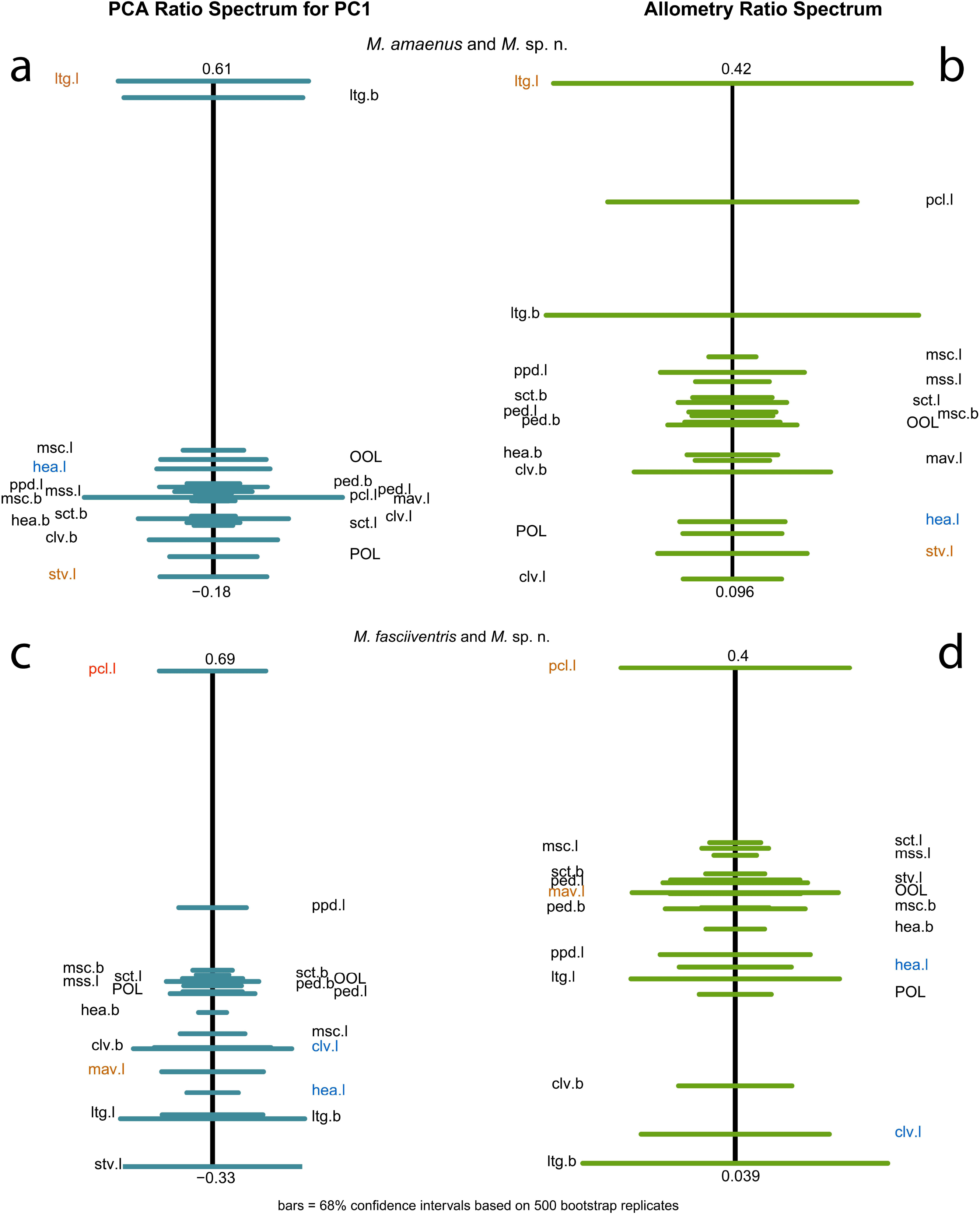
Ratio spectra for the two species pairs: a) and b) show *M. amaenus* and *Mesopolobus* sp. n, while c) and d) show *M. fasciiventris* and *Mesopolobus* sp. n. The left figures a) and c) show the PCA ratio spectrum, while the ones on right side b) and d) show the allometry ratio spectrum; horizontal bars in the ratio spectra represent 68% bootstrap confidence intervals based on 1000 replicates.

For the species pair *M. amaenus* and *Mesopolobus* sp. n. the LDA ratio extractor identified stv.l/lgt.l and hea.l/stv.l as the first two best discriminating ratios. These two combined ratios successfully separated the two species (Figure 3a). The calculated ratios for the LDA-suggested characters (*M. amaenus* vs *Mesopolobus* sp. n., range, mean, sd) are: stv.l/lgt.l (1.38-4.25, 2.38, 1.21) vs (0.80-1.15, 0.92, 0.11); hea.l/stv.l (0.11-1.12, 0.93, 0.13) vs (1.11-1.53, 1.26, 0.11). This suggests that the two species can be separated when judgement is based on a series of individuals. Further, the calculated D.shape is much higher than D.size in all of the two best discriminative ratios, indicating that species are mostly separated by differences in shape of characters (Table 3).

**Figure 3.**
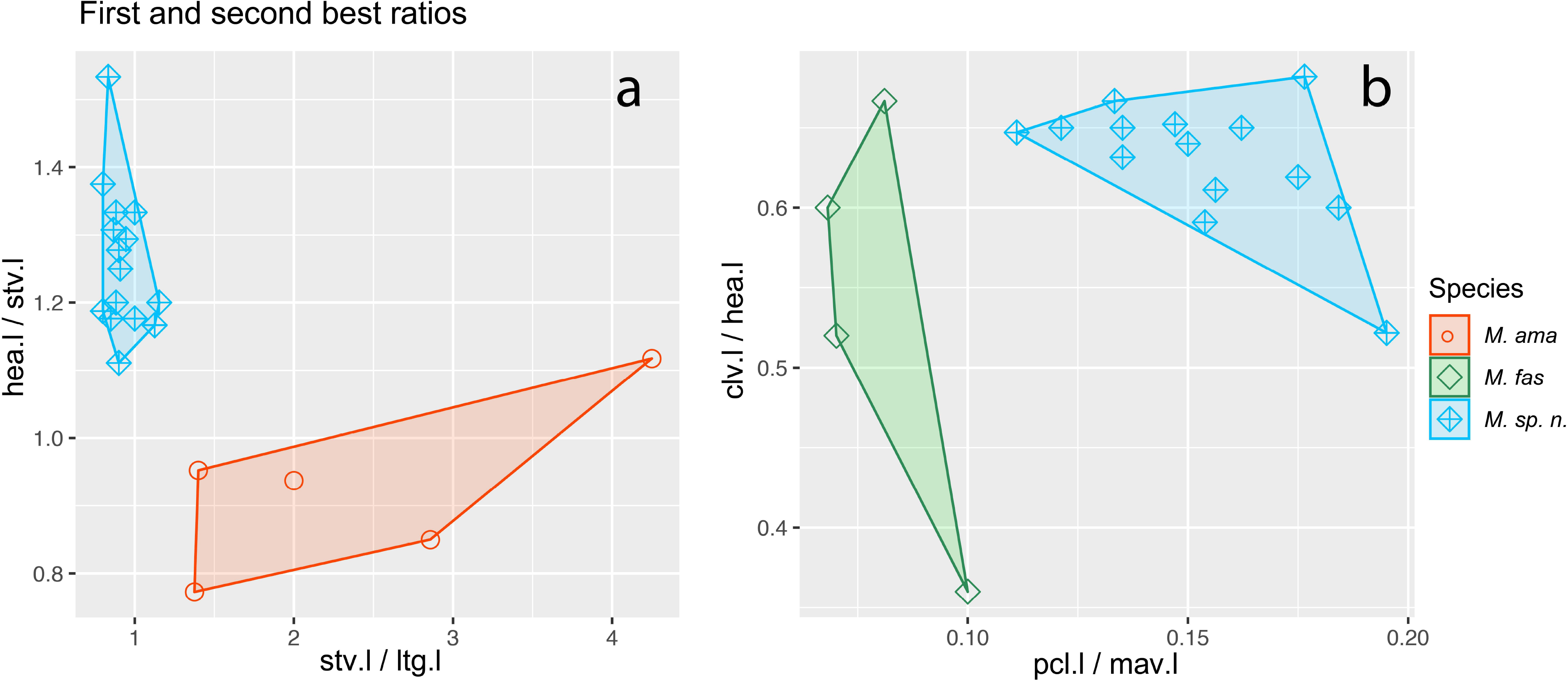
Scatterplots of the two most discriminating ratios for females a) of *M. amaenus* and *Mesopolobus* sp. n and b) *M. fasciiventris* and *Mesopolobus* sp. n. Both plots show first versus second ratio from LDA ratio extract analysis.

**Table 3.** First and second-best ratios found by the LDA ratio extractor for separating various groups and specimens of *Mesopolobus* females.

|  | Best ratios | Range group 1 | Range group 2 | D.shape | D.size |
| --- | --- | --- | --- | --- | --- |
| Group / species comparison |  |  |  |  |  |
| *M. amaenus* vs. *Mesopolobus* sp. n. | stv.l/lgt.l | 1.38-4.25 | 0.80-1.15 | 0.73 | 0.03 |
|  | hea.l/stv.l | 0.11-1.12 | 1.11-1.53 | 0.72 | 0.04 |
| *M. verditer* | stv.l/lgt.l | 1.91-2.50 |  |  |  |
|  | hea.l/stv.l | 1.14 |  |  |  |
| *M. sericeus* | stv.l/lgt.l | 1.41 |  |  |  |
|  | hea.l/stv.l | 1.1 5 |  |  |  |
| *M. fasciiventris* vs. *Mesopolobus* sp. n. | pcl.l/mav.l | 0.07-0.10 | 0.11-0.20 | 0.65 | 0.01 |
|  | clv.l/hea.l | 0.36-0.66 | 0.52-0.68 | 0.63 | 0.01 |
| *M. typographi* | pcl.l/mav.l | 0.1 |  |  |  |
|  | clv.l/hea.l | 0.55 |  |  |  |

For the species pair *M. fasciiventris* and *Mesopolobus* sp. n. the LDA ratio extractor identified pcl.l/mav.l and clv.l/hea.l as the first two best discriminating ratios. These two combined ratios successfully separated the two species (Figure 3b). The calculated ratios for the LDA-suggested characters (*M. fasciiventris* vs *Mesopolobus* sp. n., range, mean, sd) are: pcl.l3/mav.l (0.07-0.10, 0.08, 0.02) vs (0.11-0.20, 0.15, 0.05); clv.l/hea.l (0.36-0.66, 0.50, 0.14) vs (0.52-0.68, 0.63, 0.04). This suggests that the two species can be separated when judgement is based on a series of individuals. Further, the calculated D.shape is much higher than D.size in all of the two best discriminative ratios, indicating that species are mostly separated by differences in shape of characters (Table 3).

Because *M. verditer* and *M. sericeus* specimens were overlapping in the shape PCA with *M. amaenus* we calculated the best ratios for discriminating *M. amaenus* from *Mesopolobus* sp. n. for these two species as well. *M. typographi* overlapped in the shape PCA with *M. fasciiventris*, so we also calculated the best ratios discriminating *M. fasciiventris* from *Mesopolobus* sp. n. for *M. typographi* (Table 3).

### Molecular species delimitation

Based on molecular analysis of the available samples, *M. robiniae* sp. n. is placed closest to *Mesopolobus verditer* (Norton, 1868) (Figure 4). The three individuals represented only one haplotype (653 bp) that was deposited in GenBank with the MF098549 accession number. The alignment of the downloaded sequences was 468 bp long and consisted of 3 *Pteromalus* species (used as outgroup) and 14 *Mesopolobus* species, including the one described in this paper.

**Figure 4.**
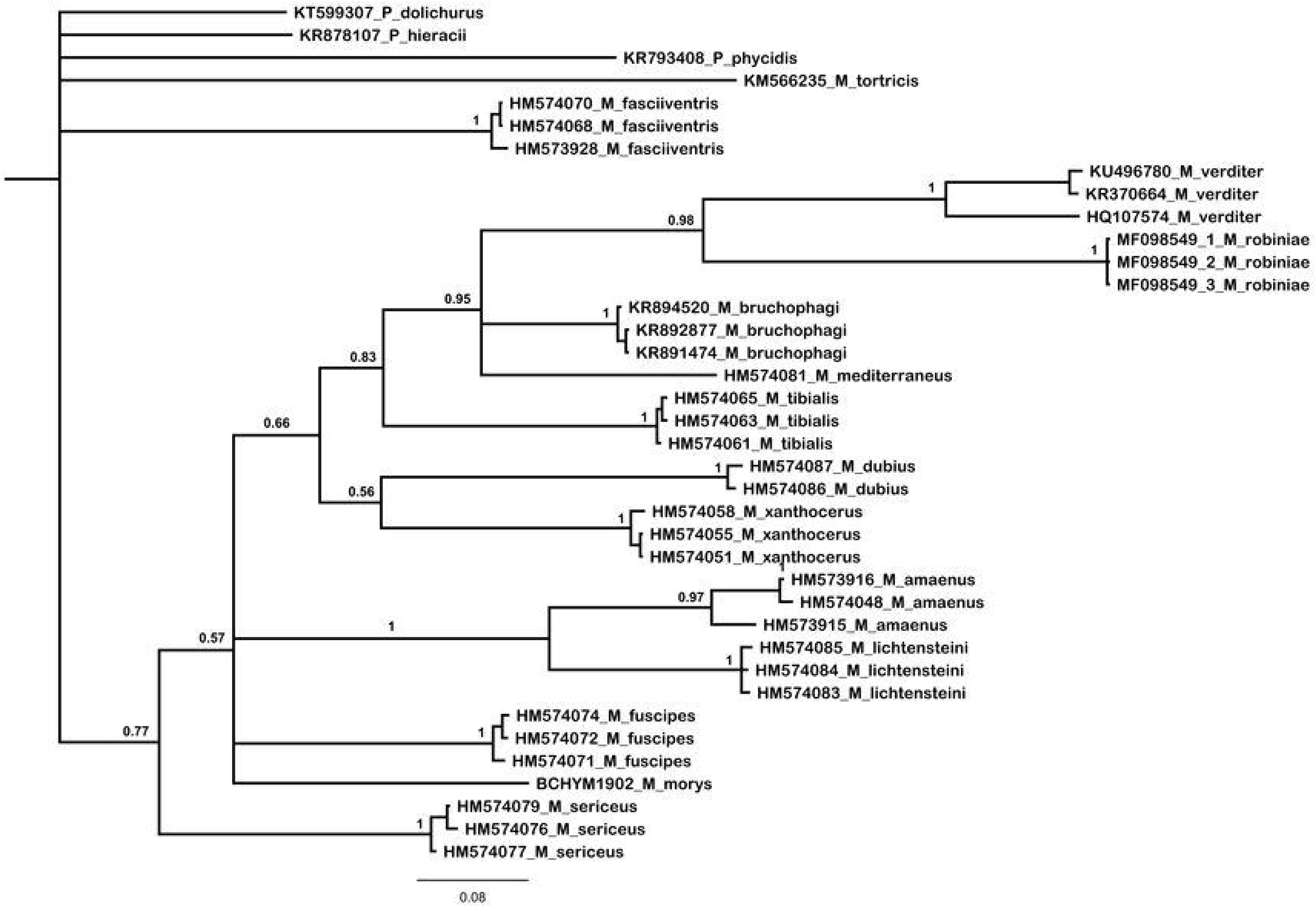
Bayesian inference (BI) tree of the *Mesopolobus* species that have available mitochondrial COI sequences. Numbers on the branches represent posterior probabilities (PP).

**Figure 5.**
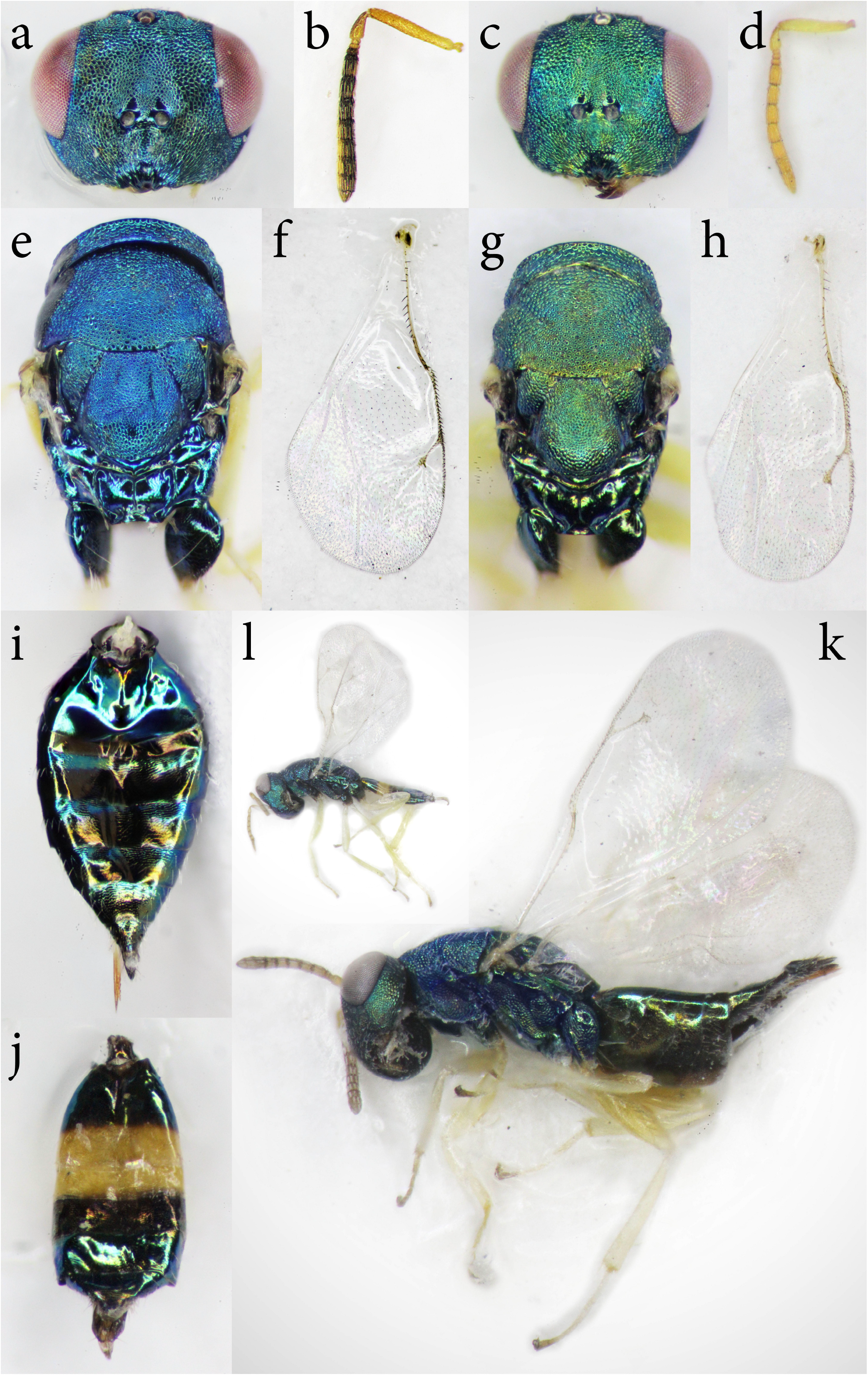
a) female head, frontal view b) female antenna c) male head, frontal view d) male antenna e) female mesosoma f) female fore wing g) male mesosoma h) male fore wing i) female gaster, dorsal view j) male gaster, dorsal view k) female habitus l) male habitus of *Mesopolobus robiniae* sp. n.

The phylogenetic relationship between the species is unresolved based on the available COI sequence data, but the tree shows a well-supported differentiation (PP=1) of the new species, with *M. verditer* as the closest species (Figure 4). The differentiation is also supported by the *p*-distance values with a minimum of 12.5% between *M. robiniae* sp. n. and *M. verditer*, and a maximum of 16.52% between *M. robiniae* sp. n. and *M. tibialis* (Table 4).

**Table 4.** *P*-distance values for sequences of *M. robiniae* sp. n. and all available *Mesopolobus* species.

|  |  |  |  |  |  |  |  |  |  |  |  |  |
| --- | --- | --- | --- | --- | --- | --- | --- | --- | --- | --- | --- | --- |
| *M. verditer* |  |  |  |  |  |  |  |  |  |  |  |  |
| *M. bruchophagi* | 12.66 |  |  |  |  |  |  |  |  |  |  |  |
| *M. amaenus* | 12.71 | 12.46 |  |  |  |  |  |  |  |  |  |  |
| *M. tortricis* | 14.22 | 13.08 | 14.98 |  |  |  |  |  |  |  |  |  |
| *M. dubius* | 15.41 | 13.54 | 14.89 | 17.66 |  |  |  |  |  |  |  |  |
| *M. lichtensteini* | 13.47 | 13.68 | 10.06 | 15.98 | 14.02 |  |  |  |  |  |  |  |
| *M. sericeus* | 14.75 | 13.87 | 14.46 | 15.6 | 15.81 | 14.45 |  |  |  |  |  |  |
| *M. fuscipes* | 13.22 | 12.39 | 13.87 | 15.75 | 13.89 | 13.16 | 11.82 |  |  |  |  |  |
| *M. fasciiventris* | 16.98 | 14.73 | 14.96 | 15.58 | 15.15 | 15.1 | 14.65 | 16.65 |  |  |  |  |
| *M. tibialis* | 15.46 | 11.2 | 15.46 | 16.82 | 14.39 | 15.74 | 14.81 | 14.25 | 16.08 |  |  |  |
| *M. xanthocerus* | 15.51 | 12.4 | 13.44 | 17.28 | 12.18 | 14.74 | 12.89 | 11.54 | 15.08 | 12.11 |  |  |
| *M. morys* | 12.68 | 10.48 | 12.76 | 14.18 | 13.66 | 13.15 | 13.9 | 10.89 | 14.69 | 13.9 | 12.03 |  |
| ***M. robiniae* sp. n.** | 12.5 | 13.54 | 14.39 | 14.68 | 14.21 | 15.02 | 14.39 | 15.81 | 15.94 | 16.52 | 15.17 | 15.85 |

### Taxonomy

*Mesopolobus* Westwood, 1833

Westwood, 1833, Philosophical Magazine (3) 2:443

Type species: *Mesopolobus fasciiventris* Westwood, by monotypy

*Mesopolobus robiniae* Lakatos & László, sp. n.

Figure 1. Female: 1a-b, e-f, i, k. Male: 1c-d, g-h, j, l.

Material examined:

Holotype, ♀, collected on 11.03.2015 near Săldăbagiu de Munte, Bihor County, 47.096354°N 21.984963°E by Lakatos, T. K, emerged on 01.04.2015, deposited in MZBBU id: MZBBU HYM000011.

Paratypes, 1♂, collected on 11.03.2015 near Săldăbagiu de Munte, Bihor County, 47.096354°N 21.984963°E by Lakatos, T. K, emerged on 01.04.2015, deposited in MZBBU id: MZBBU HYM000012; 1♀, collected on 17.03.2015 near Cluj-Napoca, Cluj County, 46.777109°N 23.674495°E by Lakatos, T. K, emerged on 10.04.2015, MZBBU id: MZBBU HYM000013; 1♀, collected on 18.03.2014 near Cluj-Napoca, Cluj County, 46.777109°N 23.674495°E by Lakatos, T. K, emerged on 05.2014, MZBBU id: MZBBU HYM000014; 2♀, collected on 17.03.2009 in Cluj-Napoca, Cluj County, 46.768086°N 23.568935°E by Lakatos, T. K, emerged on 04.2009, MZBBU id: MZBBU HYM000015 and HYM000016; 1♀, collected on 08.03.2014 near Săldăbagiu de Munte, Bihor County, 47.100895°N E21.967509°E by Lakatos, T. K, emerged on 22.04.2014, MZBBU id: MZBBU HYM000017; 1♀, collected on 11.03.2014 near Săldăbagiu de Munte, Bihor County, 47.098182°N E21.975352°E by Lakatos, T. K, emerged on 23.04.2014, MZBBU id: MZBBU HYM000018; 2♂, collected on 08.03.2014 near Săldăbagiu de Munte, Bihor County, 47.098182°N 21.975352°E by Lakatos, T. K, emerged on 22.04.2014, MZBBU id: MZBBU HYM000019 and MZBBU HYM000020; 1♂, collected on 14.03.2009 near Săldăbagiu de Munte, Bihor County, 47.079446°N 21.970817°E by Lakatos, T. K, emerged on 05.2009, MZBBU id: MZBBU HYM000021; 1♂, collected on 11.03.2015 near Săldăbagiu de Munte, Bihor County, 47.098519°N 21.984808°E by Lakatos, T. K, emerged on 04.2015, MZBBU id: MZBBU HYM000022; 1♂, collected on 02.03.2015 near Debrecen, Hajdú-Bihar County, Hungary, 47.554773°N 21.591610°E by Lakatos, T. K, emerged on 04.2015, MZBBU id: MZBBU HYM000023; 1♂, collected on 22.03.2014 near Cluj-Napoca, Cluj County, 46.834976°N 23.651004°E by Lakatos, T. K, emerged on 05.2014, MZBBU id: MZBBU HYM000024; 1♂, collected on 17.03.2009 in Cluj-Napoca, Cluj County, 46.768086°N 23.568935°E by Lakatos, T. K, emerged on 05.2009, MZBBU id: MZBBU HYM000025.

The specimens used for the genetic analysis were collected on 13.03.2014 near Săldăbagiu de Munte, Bihor County, 47.0968°N 21.98525°E and emerged on 17.04.2014.

#### Description of *Mesopolobus robiniae* Lakatos & László, sp. n

FEMALE. Length 2.05 to 3.00 mm (N=15, mean=2.6, sd=0.29 mm).

Coloration. Body green, sometimes with golden reflections; gaster bronze-black distally, some of the tergites occasionally with blue or violet flecks. Coloration of antennae: scape, pedicellus and anelli testaceous, sometimes last anellus infuscate, all funicular segments and clava always infuscate, occasionally brown. Coxae concolorous with the thorax, femora and tibiae testaceous, the tips of the fifth tarsi fuscous to black. Tegulae hyaline, usually slightly yellow posteriorly. Wings hyaline; venation pale yellow.

Head 1.1 (range 1.02-1.18) times as broad as mesoscutum; in dorsal view 2.25 (2.07-2.52) times as broad as long, with temples rounded off and between one third and one fourth as long as eyes; POL 2.11 (1.75-2.80) times OOL. Head in front view suboval with the genae moderately buccate. Eyes separated about 1.59 (1.18-1.74) times their length. Malar space more than half (0.68 (0.55-0.76)) the length of an eye. Breadth of oral fossa 1.93 (1.69-2.36) times the malar space. Clypeus strigose, its anterior margin moderately emarginate. Head uniformly and moderately reticulate. Antennae inserted low on head, lower edge of toruli at or hardly above level of ventral edge of eyes; distance between clypeal margin and toruli 0.69 (0.54-0.8) times the distance between median ocellus and toruli. Scape length 1.23 (1.09-1.4) times eye length, scape almost reaching lower edge of median ocellus; combined length of pedicellus and flagellum 0.87 (0.76-0.96) times breadth of head; pedicellus (profile) 2.06 (0.75-2.5) times long as broad, about as long as anelli plus first funicular segment; flagellum rather weakly clavate, proximally as stout as or slightly stouter than the pedicellus; first and second anelli short, twice or rather more than twice as broad as long, third anellus longer and about 1.5 times as broad as long; funicular segments subquadrate, the proximal ones sometimes slightly longer than broad, the distal ones occasionally very slightly transverse; clava 1.9 (1.5-2.29) times long as broad, 0.83 (0.66-1.15) as long as the three preceding funicular segments together; sensilla in one row on each segment, sparse on the funicle, more numerous on the clava.

Mesosoma 1.52 (1.38-1.74) times as long as broad. Pronotal collar moderately long medially, 0.21 (0.16-0.26) (one sixth to one fifth) as long as mesoscutum, and much longer at the sides, strongly and coarsely reticulate, clearly margined. Mesoscutum 1.58 (1.28-1.82) times as broad as long, rather coarsely reticulate discally, more finely laterally, without piliferous punctures. Scutellum 0.9 (0.82-0.94) as broad as long, moderately convex, finely reticulate, the frenum rather more coarsely reticulate. Axillae finely reticulate. Dorsellum a narrow, alutaceous transverse crest which is separated from the scutellum by a simple suture. Propodeum medially slightly less than half (0.41 (0.36-0.48)) as long as the scutellum; median area 2.39 (2-3) times as broad as long, well-defined laterally, the plicae distinct throughout and sharp over at least their distal half; median carina distinct, straight; panels of median area finely, slightly irregularly reticulate; nucha transversely aciculate, separated from the median area by an impressed line; posterior foveae, at sides of nucha, moderately deep; spiracles oval, longer than broad, separated by nearly half their length from the metanotum. Postspiracular sclerite broad, shiny, weakly and irregularly sculptured. Mesepisternum moderately finely reticulate, its upper triangular area smooth; mesepimeron rather more coarsely reticulate than the mesepisternum, metapleuron smooth. Legs rather short; femora rather stout; mid tibiae fairly slender, 7.44 (4.88-9) as long as their maximum breadth. Fore wing rather broad; costal cell fairly broad, its upper surface bare, lower surface with a complete row of hairs and some additional hairs scattered over the distal third to half; basal cell bare, open below; basal vein bare or with one to three hairs; speculum open below, on upper surface of wing extending below the proximal end of the marginal vein; surface beyond the speculum thickly pilose; marginal vein 2.19 (2-2.47) times as long as the stigmal vein; postmarginal vein shorter than the marginal, 0.73 (0.63-0.81) times as long as the marginal.

Gaster ovate, 1.24 (1.16-1.33) times longer than mesosoma, 0.8 (0.66-0.96) times broader than mesosoma, 2.37 (1.91-2.96) times as long as broad; basal tergite occupying from slightly more than one quarter, to nearly one third, the total length; last tergite somewhat shorter than its basal breadth, its length 1.07 (0.72-1.79) times its breadth; ovipositor sheaths projecting at most very slightly; hypopygium slightly reaching the middle of the gaster, ratio of hypopygium length to gaster length is 0.44 (0.35-0.54).

MALE. Length 1.8 to 2.25 mm (N=15, mean=2.02, sd=0.16 mm).

Coloration: head and mesosoma bright green; gaster greenish dorsally with T2 and posterior half of TI yellow, T3 purplish; antennae bright testaceous; legs except coxae yellow, last tarsal segments grey-brown.

Head: antenna with 3 anelli and 5 funicular segments, length of pedicel plus flagellum 0.97 (0.85-1.04) times breadth of head; scape 5.33 (4.6-6.25) times as long as broad, without a boss on its anterior surface; flagellum proximally not broader than pedicel, F1-F4 longer than broad, F5 subquadrate. Mouthparts unmodified; no patch of modified sculpture behind malar sulcus.

Mesosoma: Pronotal collar long as in females, about 0.21 (0.18-0.27) mesoscutal length. Middle tibiae unmodified, middle tibia 7.42 (6.29-8.8) times its breadth, tibial spur 1.47 (1.17-1.8) times breadth of first tarsus.

Gaster oblong, ovate, 0.91 (0.83-1.02) times as long as mesosoma, 2.24 (1.63-2.63) times as long as broad with a yellow ventral plica; T1 with triangular depression at base.

### Etymology

The new *Mesopolobus* species is named after the host plant of its seed predator host, the black locust (*Robinia pseudoacacia*).

#### Diagnosis

##### Morphological comparison

*M. robiniae* sp. n. females were not identifiable based on Graham’s keys (Graham, 1969), but several morphologically and morphometrically related species were found for which the differing characters will be enumerated in the order that the species appear in Graham’s key. The species *M. robiniae* sp. n. has a shorter gaster compared to head plus thorax than *M. maculicornis*. The species *M. jucundus* has a curved stigmal vein compared to *M. robiniae* sp. n. *Mesopolobus robiniae* sp. n. differs from *M. fasciiventris* by its males having 3 anelli and 5 funicular segments while in the latter there are 2 anelli and 6 segments. The head of *M. apicalis* in dorsal view has temples nearly three quarters as long as the eyes, while *M. robiniae* sp. n. has its head in dorsal view temples appearing one quarter to one third as long as the eyes. The gaster of *M. amaenus* is less than twice as long as its breadth and almost as long as the thorax, while in the case of *M. robiniae* sp. n gaster is not less than twice as long as broad, but it is as long as thorax. The species *M. longicollis* has the pronotal collar 1/7 to 1/6 as long as the mesoscutum and its gaster is less than twice as long as broad compared to *M. robiniae* sp. n. The species *M. diffinis* and *M. meditteraneus* differ from *M. robiniae* sp. n. because the latter has longer marginal vein as 1.4 to 1.6 as length of the stigmal vein.

The species *M. verditer* is not present in the keys of Graham (1969) because it has a North-American distribution. It differs from *M. robiniae* sp. n. in the following: antennal funicle segments shorter than their length, while in *M. robiniae* sp. n. they are at least as long as their breadth. The ratio of the stigma vein to the last gastral tergite length is 1.91-2.50 in *M. verditer*, while in *M. robiniae* sp. n. is between 0.08-1.15. *Mespolobus sericeus* differs from *M. robiniae* sp. n. first by having 2 anelli and 6 funicular segments, but also in having the ratio of the stigmal vein to the last gastral tergite length 1.41, while in the other species this ratio is smaller (0.8-1.15). From *M. typographi* the species *M. robiniae* sp. n. differs in the ratio of the pronotal collar length to the marginal vein length which in the first species is 0.1 (N=1) and in the second is between 0.11-0.02 (N=15). Moreover, in *M. typographi* the median area of propodeum is 1.75-2 times as broad as long (Graham 1969) while in *M. robiniae* sp. n. is 0.82-0.94 times as broad as long (N=15).

Based on von Rosen’s key (von Rosen, 1958), the morphological identification of specimens led us to *M. mediterraneus* (Mayr, 1903) as the closest species, from which *M. robiniae* sp. n. females differed in having a longer pronotal collar, much longer marginal than stigmal vein and a shorter gaster than the combined length of head and mesosoma.

In Gahan (1932) page 739 says that *Mesopolobus* (syn. *Amblymerus*) *verditer* (Norton, 1868) “… conforms very closely to the characters of the genus *Amblymerus* Walker as represented by *Amblymerus amoenus* Walker…” (syn. *M. amaenus*), when transferring the species to genus *Amblymerus* Walker, 1834 from the genus *Nasonia* Ashmead, 1904. The hosts of *M. verditer* are usually sawflies (Hymenoptera: Diprionidae) on pines (*Pinus* sp.) (Noyes, 2020). *M. verditer* is distributed in the Nearctic and Germany (W. R. Thompson, 1958). Moreover, *M. verditer* differs from *M. robiniae* sp. n. in having a reticulated middle area of propodeum and oblique wrinkles, as does also from *M. amaenus* and *M. longicollis* (von Rosen, 1958).

We propose the following update to the key of *Mesopolobus* species of Graham (1969) for females:

> 16(14) - Either gaster at least slightly longer than head plus thorax, and usually more than twice as long as broad, or gaster not longer than head plus thorax, and at most twice as long as broad.……………………………………….…………………………………………….…….. 16A
>
> – Gaster not longer than head plus thorax, their ratio is 0.94 (0.88-0.98), gaster usually more than twice, 2.37 (1.91-2.96) as long as broad …………………………….. *M. robiniae* sp. n.
>
> 16A(16)- Gaster at least slightly longer than head plus thorax, usually more than twice as long as broad ……………………………………………………………………………………….. 17
>
> – Gaster not longer than head plus thorax, at most twice as long as broad …………………. 27

##### Distribution

The type locality for *M. robiniae* sp. n. is Săldăbagiu de Munte, Bihor County, Romania (N47.096354 E21.984963). The other localities are situated in the neighbouring counties: Cluj County, Romania and Hajdú-Bihar County, Hungary. The species may appear in the Carpathian Basin where its host plant is present, but we expect that it may also be found outside of the Carpathian Basin, in Eastern Europe and maybe throughout Europe.

### Biology

Based on our rearing, *M. robiniae* sp. n. seems to be an early flying parasitoid species. Individuals of the species emerged during spring consequently in all study years. Our black locust seedpod samples were collected mostly in March, and the peak of *M. robiniae* emergence was in April, with a decrease in May. After May we rarely encountered any individuals of this parasitoid species.

The host of *M. robiniae* sp. n. may be *Bruchophagus robiniae* but there is no information regarding the host plant of *B. robiniae* before the introduction of black locust. Another possibility is that *M. robiniae* sp. n. initially had another host, but has switched from it to *B. robiniae*. Either possibility is plausible; before 1970 (Zerova, 1970) the species *B. robiniae* was not known, and *M. robiniae* sp. n. was not described until now. The parasitoid community of black locust is understudied, and the available literature makes no mention of parasitoids in this community (Farkas & Terpó-Pomogyi, 1974; Perju, 1998), with the exception of our ecological study concerning the seed-predator community of black locust in Eastern Europe (Lakatos *et al*., 2016).

## Discussion

The multivariate ratio analysis (MRA) and the *mt*DNA sequence analysis resulted in the successful separation of the *Mesopolobus* species emerging from black locust seedpods from the other congeneric relatives. The morphometry-based shape PCA helped us identify which species fall closer to the specimens emerged from black locust crops. This delimitation was important since the available specific keys (Graham, 1969) did not lead us to a closest relative based on the combination of morphology and morphometry.

In a PCA ratio spectrum, only ratios calculated with variables lying at the opposite ends of the spectrum are relevant for a particular shape PC and the most allometric ratios are also found at the opposite ends of the allometry ratio spectrum (Baur *et al*., 2014). The PCA ratio spectrum and the allometry ratio spectrum plots revealed a large (*M. amaenus* and *M. robiniae* sp. n.) and moderate (*M. fasciiventris* and *M. robiniae* sp. n.) amount of allometric variation in the identified discriminating morphometric character pairs (Figure 2). However, this is not of concern in our case, because on one hand the species we found to be closely related based on morphometry were clearly separated based on the molecular results, and the combinations of the usually used ratios do not overlap with species in the keys of Graham, since there is no possibility to progress beyond key couplet 16. On the other hand, the ratios found with the LDA ratio extractor tool have small D.size values compared to D.shape values (Table 3) which means that separation was mainly due to shape rather than size. The LDA ratio extractor tool found that the species pairs could be separated without overlapping based on the first ratio pairs. These ratios in combination with morphologic characters gave a confident separation of the closely related species.

The origin of *M. robiniae* sp. n. species is yet unknown, since it has to be a host shifting species. Black locust was introduced to Europe 300 years ago, and in its native area it has no *Bruchophagus* seed predator, nor the associated parasitoids (Stone, 2009). So, the new *Mesopolobus* species may not be monophagous on *B. robiniae*, which is similarly a host-shifting seed predator. Nonetheless, it is befitting of the name *robiniae*, since parasitoids are also affected by the host plant of their herbivorous host. As part of their host finding strategy, parasitoids may search for a specific plant or plant part (as seedpods) housing any potential herbivorous host species (Cronin & Abrahamson, 2001).

## Acknowledgements

We thank to Zoltán Vas, Curator of Hymenoptera collection, Hungarian Natural History Museum for loaning several specimens of various *Mesopolobus* species and for his valuable help during identification and manuscript preparation. We are thankful to Natalie Dale-Skey, curator of the Hymenoptera section, Natural History Museum for loaning several *Mesopolobus* specimens and to James Hogan Collections Manager of Hope Entomological Collections, Oxford University Museum of Natural History for providing photography of *M. longicollis*. We are also thankful to Lajos Király for his help in the molecular analysis. The authors are grateful to Chris Looney (Washington State Department of Agriculture, Olympia, United States) for his review, comments and suggestions of the manuscript. Molecular analysis was done at the Interdisciplinary Research Institute on Bio– Nano–Sciences of BBU, Cluj, Romania. During preparation of the manuscript AL Dénes received financial support from the Collegium Talentum scholarships, Hungary.

## Author contributions

LZ and LKT designed the study. LKT collected data, made morphometric measurements, participated in paper writing. DAL made molecular analyses, participated in paper writing. LZ analyzed, interpreted data and drafted the manuscript. All authors gave final approval for publication.

